# Swine reporter model for preclinical evaluation and characterization of gene delivery vectors

**DOI:** 10.1101/2025.06.13.659546

**Authors:** Jarryd M Campbell, Derek M Korpela, Hesong Han, Sheng Zhao, Dennis A Webster, Yen Anh H Nguyen, Robyn O Aune, Hinsoukpo Dagan, Rebecca Milliken, Jonathan K Watts, Niren Murthy, Daniel F Carlson

## Abstract

Delivery of gene therapy vectors targeted to any somatic cell remains a key barrier for the development of genetic medicines. While rodent models provide insights into vector biodistribution and cellular tropism, their anatomical and physiological differences from humans limit their translational potential and studies in large animal models are often required. In this study, we developed a swine reporter model (SRM-1) to evaluate both viral and nonviral vector delivery in a large animal system. The SRM-1 model harbors a Cre- and CRISPR-activated tdTomato reporter at the *Rosa26* locus that allows for tracing of cell-specific delivery and expression of gene therapy vectors *in vivo*. To evaluate this model, we administered adeno-associated virus serotype 9 (AAV9) and lipid nanoparticles (LNPs) carrying mRNA systemically and found successful *in vivo* reporter activation across a variety of tissues. Intracerebroventricular (ICV) administration of LNP-Cre mRNA was also performed and demonstrated localized activation in cortical brain cells. In addition to biodistribution studies, this model has utility for testing safety and clinically relevant administration methods, surgical and non-surgical, of delivery vectors. Our findings support the SRM-1 model as a valuable tool for advancing gene therapies from preclinical testing to clinical application.

## INTRODUCTION

Safe and effective targeting of somatic cells with viral or nonviral vectors is a continuous barrier to gene therapy development. These vectors can be used to deliver payloads of therapeutic nucleic acids or proteins to treat diseases, especially in those with genetic underpinnings^1^. Rodent models have been essential in characterizing the tissue and cellular tropism of these viral and nonviral payloads^2–7^. Fluorescent reporter mice, activated by genome-editing enzymes like Cre or CRISPR-Cas9, are critical to help track tissue and cell uptake of delivery vectors and payload deployment *in vivo*. For example, reporter mice have aided in the characterization of lipid nanoparticle (LNP)-based mRNA delivery for *in vivo* gene delivery, with several formulations demonstrating both hepatic and extrahepatic tropism when delivered systemically^4–6,8,9^. However, anatomical and physiological differences between rodents and humans create limitations when translating these discoveries to humans. This is especially true when studying the impact of alternate routes of administration, dosing in large animals, and the role of the immune system in vector targeting and expression. At advanced stages of preclinical testing, nonhuman primates (NHPs) have been used to study vector delivery and safety *in vivo* as they are more physiologically like humans. However, their small size, high cost and low fecundity limit utility and access for most researchers. Alternative preclinical animal models, especially large animals that are translationally relevant, can be useful in exploring novel vector and gene delivery to tissues and cells.

In this study, we have developed a swine reporter model as part of the NIH’s Somatic Cell Genome Editing Consortium for characterization of both viral and nonviral vectors in a more accessible large animal model^10^. Swine are an increasingly attractive species for preclinical studies and have several advantages for use in biomedical research. First, the technology development around genome engineering and reproductive technologies has matured and custom model development is readily accessible. Highlighting these efforts are the recent advances in xenotransplantation of multi-edited swine organs^11^, which demonstrates the utility of these technologies that are also being used to generate novel pig models for studying disease^12^. Pigs are available in a variety of body sizes to accommodate a wide range of gene delivery studies, from systemic infusions to local delivery with medical devices. Shared physiological and anatomical characteristics exist between pigs and humans, particularly in the nervous system^13^, the cardiovascular system^14^, the gastrointestinal tract^15^, the respiratory system^16^, the eye^17^, and skin^18^. Finally, swine husbandry and breeding are efficient due to the centuries-old reliance of pigs as a major food source. Pigs have large litters compared with other large animal model systems, typically rearing 5-15 piglets per litter depending on the breed. These piglets develop rapidly to reach sexual maturity in 6 months, and gestation lasts only 4 months. This rapid cycling through generations makes it possible to run cost efficient, translationally relevant, and more robust animal studies comparted to NHP.

## RESULTS

### Generation of the SRM-1 porcine model

We designed a swine reporter model (SRM-1) based on the Ai9/Ai4 mouse reporter^2^. It harbors a constituently active CAG promoter that drives expression of a tdTomato fluorescent protein after removal of a strong transcriptional stop domain (3xStop, Fig. 1A). The 3xStop is flanked by loxP sites, which can be removed by Cre recombinase, as well as SpCas9 and AsCas12a guide RNA spacer sequences for one-guide genome editing activation (Fig. 1A). In addition to the tdTomato reporter, a sodium iodide symporter protein is activated allowing for real-time imaging of reporter activation using PET or SPECT imaging in-life^19^. The SRM-1 reporter construct was integrated at the *Rosa26* gene using homology directed repair in Yorkshire swine fibroblasts and cloned by somatic cell nuclear transfer to yield a founder breeding population and fetal fibroblasts (Fig. S1). Resulting animals were screened for precise reporter integration by junction PCR of the SRM-1 construct into the *Rosa26* gene at the 5’ and 3’ end of the integration site (Fig. 1B). These founder animals harbored two copies of the reporter (Fig. 1B), and homozygosity of the reporter allele was confirmed by outcrossing to wild type animals to yield all heterozygous offspring (data not shown). tdTomato expression is apparent in SRM-1 fetal fibroblasts after transfection of either Cre mRNA or CRISPR ribonucleoproteins (Fig. 1C).

**Fig. 1.**
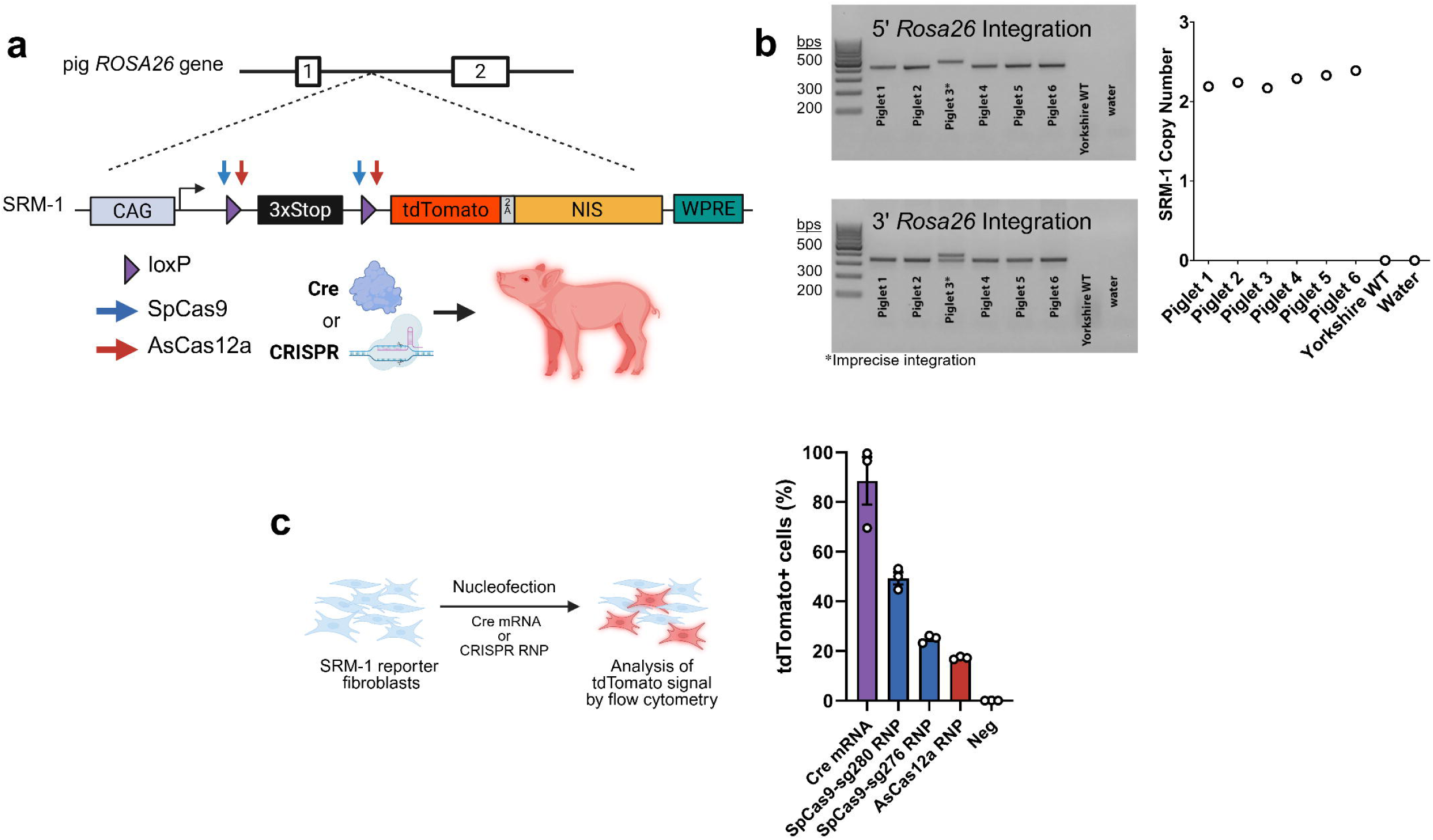
Generation of the SRM-1 Cre- or CRISPR-inducible swine reporter model. **(A)** Schematic illustration of the **SRM-1** reporter integrated into the swine *Rosa26* gene. Reporter design includes loxP sites (purple triangles) and CRISPR sites (blue and red arrows) for Cre- or genome editing-mediated reporter activation, respectively. **(B)** PCR of the integration junction of the SRM-1 construct in the *Rosa26* gene and SRM-1 construct copy number measured by ddPCR in founder SRM-1 animals. Asterisk indicates an animal with imprecise reporter integration. **(C)** %tdTomato activation in SRM-1 fibroblasts measured by flow cytometry after transfection with either Cre mRNA, SpCas9 RNP, or AsCas12a RNP. Closed circles are individual data points, error bars are standard error of the mean.

### AAV9-Cre injection into SRM-1

To assess SRM-1 reporter activation *in vivo*, adeno-associated virus serotype 9 (AAV9) vector expressing Cre recombinase was infused systemically into a male SRM-1 piglet at a dose of 7x10^12^ genome copies per kilogram body weight (Fig 2A). A droplet digital PCR (ddPCR) assay was developed both to detect Cre-mediated DNA recombination of the SRM-1 reporter and to measure distribution of the AAV genome in each tissue. Biodistribution of the viral genome and reporter recombination rates demonstrated widespread vector tropism (Fig. 2B, Fig. S2). Four of the top nine tissues with over 5% SRM-1 recombination were muscle tissues (skeletal muscle, diaphragm, heart, tongue), an expected result based on vector tropism of AAV9 in both large and small mammals^20–22^. Concurrent immunohistochemistry performed on select tissues with both high and low reporter recombination levels demonstrated robust reporter expression (Fig. 2C, Fig. S3). A subset issues showed high viral load with minimal reporter activity, most notably in the spinal cord and adipose tissue (Fig. 2B, Fig. S2). In addition to somatic cells, it is critical to assess the potential for germ cell exposure to gene delivery vectors. Substantial activation in the testicle was observed (23%), which appear isolated to cells surrounding the seminiferous tubules (Fig 2C). These results support previous findings of effective AAV9 uptake into testicular tissue, particularly testosterone-producing Leydig cells, observed in mice and NHPs^22–24^.

**Fig. 2.**
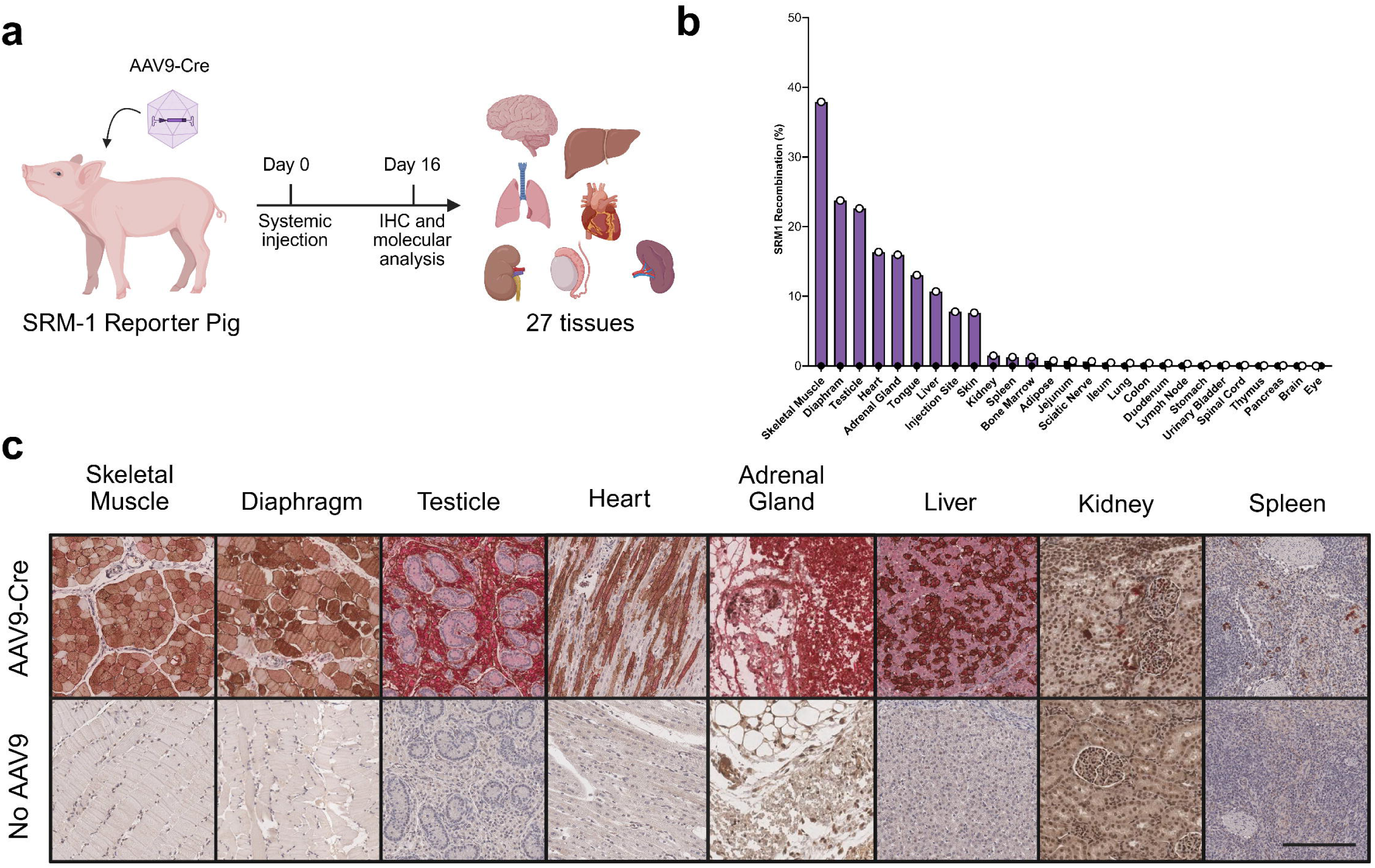
SRM-1 detects recombination *in vivo* by intravascular delivery of AAV9-Cre. **(A)** Schematic illustration of systemic AAV9 *in vivo* study. **(B)** ¾SRM-1 recombination rates of the loxP sites in tissues measured by ddPCR. Open circles (purple bars) are an AAV-injected animal (n=l), closed circles are an uninjected animal (n=l). (C) Immunohistochemistry of tissues taken from an injected and uninjected SRM-1 piglet. Tissues were co-stained for tdTomato and sodium iodide symporter (NIS) and detected with DAB. Scale bar, 300 µm.

### LNP-mRNA injection into SRM-1

Nonviral delivery of nucleic acid like mRNA is a growing therapeutic modality that has applications across multiple areas of disease treatment and management^25^. To determine the utility of the SRM-1 model for characterizing tissue-specific LNP delivery, three piglets underwent systemic infusion with LNPs^7^ at a dose of 0.4 milligrams per kilogram body weight (Fig. 3A and B, Fig. S4). Comprehensive serum chemistries and complete blood cell counts were collected on days 1, 3, 7, and 14 to analyze potential toxicities related to systemic LNP exposure (Fig 3C). No enzyme elevations were observed in animals injected with LNP, while moderate decreases in lymphocytes and increases in neutrophils and white blood cells were observed one day post injection in two LNP injected animals compared with unexposed siblings (Fig 3C). These changes in blood counts recovered to normal values by day three.

**Fig. 3.**
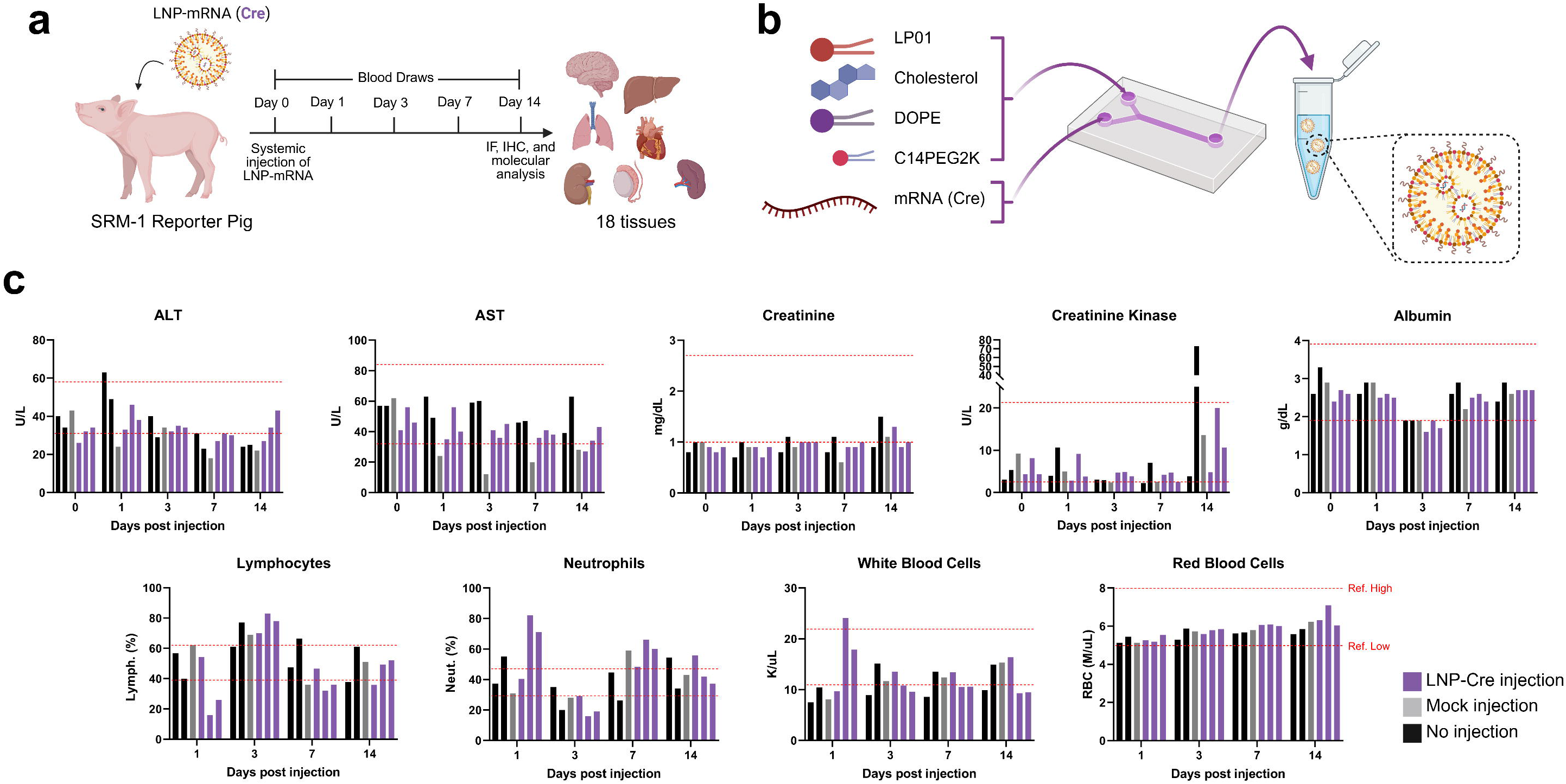
Serum chemistry and blood counts of LNP-injected piglets. **(A)** LNP-mRNA delivery study design schematic. **(B)** LNP formulation schematic of mRNA using microfluidic mixing. **(C)** Absolute values of serum ALT, AST, creatinine, creatinine kinase, and albumin as well as blood cell counts (white blood cells, lymphocytes, neutrophils, red blood cells) in three LNP-injected SRM-1 pigs (purple), one mock-injected SRM-1 pig (grey), and 2 non-injected wild type sibling piglets (black). The red dashed lines are the high and low references ranges supplied by the diagnostic laboratory. One blood sample per piglet was taken at each time point.

Of the 18 tissue types sampled, the liver had the highest DNA recombination rate (39%, Fig 4A). Samples were collected at three locations (proximal, middle, and distal from the portal vein) on each of four pig liver lobes and uniform distribution of LNP was observed throughout the tissue (Fig 4B). Body-wide tissue distribution suggests extrahepatic targeting was also achieved, with moderate rates of targeting in the spleen (26%), lung (7%), adrenal gland (5%), and measurable uptake in the upper GI tract (1-2%, Fig. 4A and C, Fig. S5). Gonad sampling from one injected male showed moderate rates of targeting to testicular cells (13%), which appear to be excluded from the seminiferous tubules. Minimal ovarian tissue uptake was observed from two injected females (1%, Fig. 4A and C, Fig. S5).

**Fig. 4.**
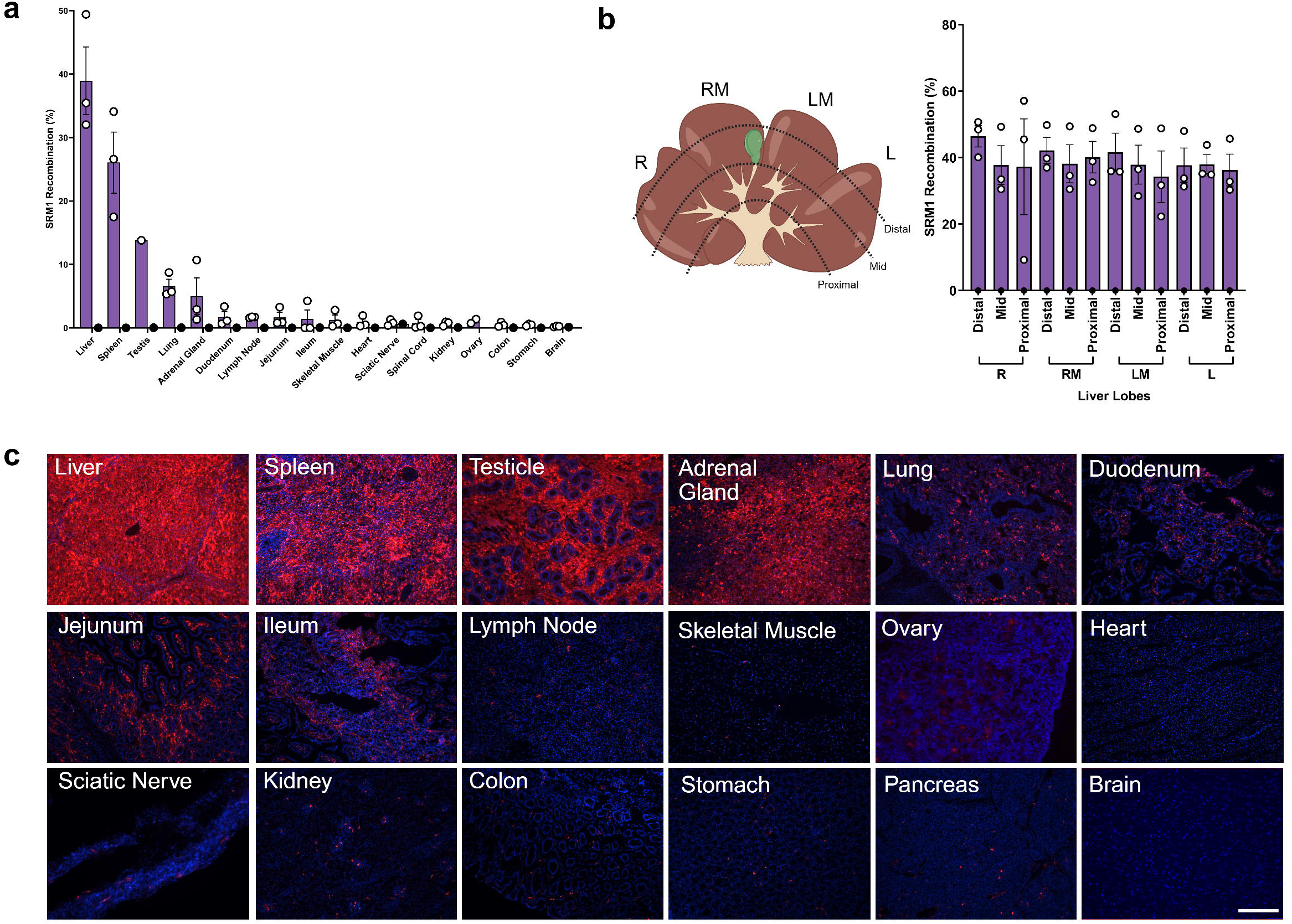
Tissue and cellular transfection by systemic LNP-mRNA can be tracked using SRM-1 swine after multiple weeks. **(A)** Recombination of SRM-1 genomic DNA collected from LNP-injected SRM-1 pigs (n=3, 1 male and 2 female) and a mock-injected male SRM-1 pig (n=l). Open circles are individual animal measurements for LNP-injected animals, closed circles are for mock-injected. Error bars are standard error of the mean. **(B)** Distribution of LNP in each swine liver lobe. R = right lateral lobe, RM= right medial lob, LM = left medial lob, L = left lateral lobe. Open circles are individual animal measurements. (C) Fluorescent histological images of SRM-1 tissues with native tdTomato-producing cells (red) and stained nuclei (blue) after LNP-mRNA (Cre) injections. Scale bar, 200 µm.

### Intracerebroventricular LNP-mRNA delivery

With the goal of achieving high levels of targeting and reducing potentially undesired physiological effects by delivering lower doses, localized delivery is an attractive alternative to systemic delivery of gene therapy vectors. Delivery of viral or nonviral vectors to the brain is one such example that holds great promise for the treatment of neurologic diseases^26^. To test reporter activity in the brain of an SRM-1 model, we performed a stereotactic injection of LNP-mRNA (Cre) into the intracerebroventricular space in a female piglet. LNP uptake was successful but limited to the outer cortex, likely due to the distribution of the vector in cerebrospinal fluid (Fig. 5A). Examination of the stained sections suggest that multiple cell types in the brain were activated along the outer cortex based on differential morphology and reporter expression patterns (Fig. 5B). Overall, this model holds promise for characterizing nonviral delivery of nucleic acid to the brain, and translatable surgical procedures can be deployed to optimize regional tissue targeting.

**Fig. 5.**
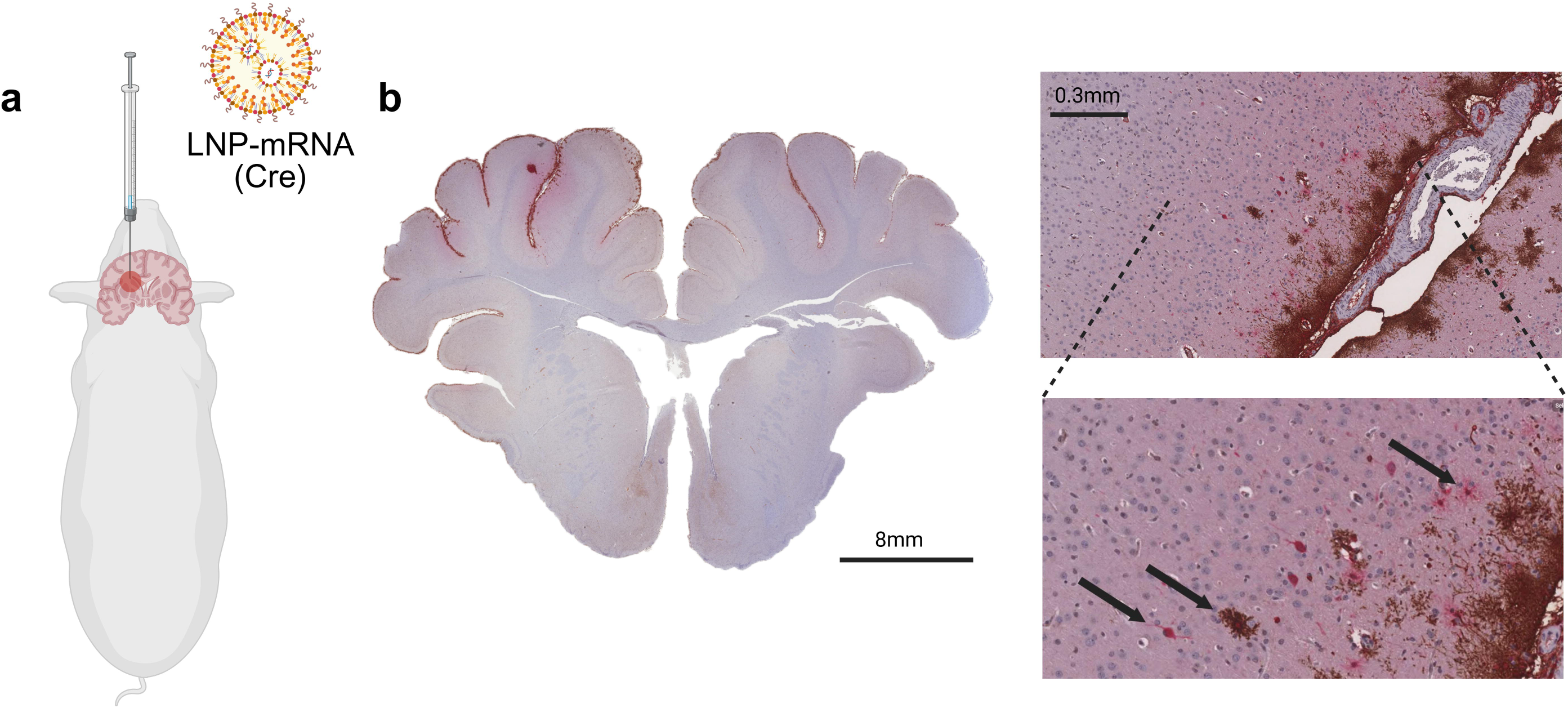
Delivery of LNP-mRNA to the brain via intracerebroventricular (ICV) injection. **(A)** Schematic of stereotactic ICV injection of LNP­ mRNA (Cre) in an SRM-1 piglet. **(B)** Whole-brain coronal sectioning and IHC for tdTomato and NIS was performed to identify mRNA uptake (left panel). Cell transfection was restricted to the outer cortex with minimal penetration into tissue. Three distinct cortical cell types of the brain were identified by morphology and staining differences (arrows, right panel).

## Discussion

Novel large animal models that provide critical insights into the complexity of vector delivery are needed to accelerate the application of gene therapies for treating a variety of diseases. Here we highlight a porcine model that can be used to track delivery of viral or nonviral vectors by activating a genetically-encoded reporter. Activation of the reporter requires vectors to traffic to the tissue, gain entry to the cells, and deliver a functional payload. We first chose to test AAV to validate the use of an established and well characterized vector in SRM-1 for *in vivo* studies. AAV9 has a wide tissue tropism in mice, which was also observed in this study^23,27^. Notably, it can cross the blood brain barrier in both NHP and mice, a discovery which translated to a gene therapy drug for spinal muscular atrophy^28,29^. Interestingly, we found vector genomes in the spinal cord but no reporter recombination or positive cell staining. AAV9 tropism in swine motor neurons has been previously described by intrathecal administration of the vector^30^, and one study in pigs also found uptake in the brain after intravenous injection at nearly twice the dose and with gene expression driven by a CMV promoter^31^. The AAV9 vector here expressed Cre from a chicken beta-actin promoter and a CMV enhancer (CB6 promoter), which has not been extensively studied in pigs but has been successfully used to cross the blood brain barrier (BBB) and express genes in the mouse brain and motor neurons along the spinal cord^32^. Our data supports no functional expression of genes in the spinal cord and little in the brain, but additional replicates, doses, and promoters should be tested to continue characterizing pigs as a preclinical model for vector studies crossing the BBB.

Unlike AAV vectors, nonviral vectors like LNPs do not have preexisting, natural extrahepatic tissue tropism and cannot localize and express their genome in the nucleus^33^. Targeting of the LNP to different cell types depends on several factors including the chemical makeup of the LNP, the delivery route to dictate which tissues are exposed, exogenous proteins that can interact with the LNP and mediate receptor targeting and uptake into the cell, and ultimately endosomal escape and intracellular trafficking of the cargo^34^. The LNP system used in the systemic study here contained the biodegradable ionizable lipid LP01 along with helper lipids, PEG-DMG, and cholesterol and has previously been shown to target and express Cas9 mRNA effectively in the liver for genome editing^7^. Replacing Cas9 mRNA with Cre mRNA, we found the highest levels of targeting in the liver and the spleen, along with testicular tissue (non-germ cell) and adrenal gland. Lung tissue showed recombination levels of the reporter at similar levels to adrenal, but tdTomato expression from fixed tissue appears to be lower. This is likely due to sampling differences between the fixed tissue and DNA collections, as the adrenal gland samples had a high amount of variance in tdTomato expression in the tissue after sectioning (data not shown). Surprisingly, low levels of reporter targeting were measured across many other tissues as well. These findings highlight the sensitivity of the model and are congruent with recent evidence that pigs express protein from LNP-based mRNA delivery even in extrahepatic tissue^35^. Little research exists detailing the tissue distribution of LNP-mRNA after systemic injection in large animals, and more in-depth studies are needed to determine the translatability of these results to humans.

The safety of systemic LNP exposure is important to study in large animals. Complement activation-related pseudoallergy (CARPA) has been observed in both human and swine in response to systemic nanoparticle exposure^36^. Because of the impact on overall health, animals in this study were monitored for a CARPA response and showed no symptoms through the procedure and in-life observations. Overall, complete blood count and serum chemistries were within normal limits throughout the study, however mild variabilities in white blood cell and neutrophil counts in two of the three LNP-injected animals was noted at 24-hours post injection which may be suggestive of a mild immune response to the LNP-mRNA vector or anesthetic procedure.

The ability to test both systemic and localized delivery through clinically relevant procedures is one of the greatest advantages swine models offer. The central nervous system can only be reached systemically by a subset of vectors that cross the BBB. Viral vectors have be reprogrammed to target novel receptors that facilitate this process, which has recently been exploited to cross the BBB using the human transferrin receptor^37^. For large nonviral molecules like LNPs this is currently not feasible, so alternative methods like intrathecal delivery, convection-enhanced tissue injection, and focused ultrasound are being explored, all of which would greatly benefit from preclinical testing in large animal models. Demonstrating this utility by performing an ICV injection, we found reporter activation in cortical brain cells when exposed to LNPs from the cerebral spinal fluid. The LNP formulation contained the ionizable lipid 306-O12B, which has been found to effectively target the liver after intravenous injection^38^ but has not been well studied for brain targeting. This formulation was not able to penetrate outer cortex tissue more than 200-300 µm from the CSF at a dose of 80 µg. LNPs will higher tissue penetration will be required to enable therapeutically relevant delivery using this method.

To keep the dosage of vector low, our studies relied on dosing juveniles (aged 3 days to 3 weeks). Systemic studies in older swine that are similar in body mass to humans will be informative to understand the impact age and size can have on overall biodistribution. As *in vivo* studies with the SRM-1 model move to gene editing-based studies, we anticipate CRISPR-based delivery will require higher doses than Cre recombinase to activate the reporter, which has been previously observed in reporter mice^6^ and through SRM-1 cell studies here. Overall, the SRM-1 swine reporter model enables precise tracking of viral and nonviral delivery and activity across multiple tissues, providing a scalable, immune-intact large animal model to advance gene delivery discovery and translation.

## Methods

### SRM-1 plasmid generation

The Ai9/Ai14 reporter sequence^2^ was modified to include SpCas9- and AsCas12a-targeting sequences flanking the 3xStop domain using Geneious Prime (Dotmatics). The resulting reporter cassette was synthesized and subcloned into a plasmid backbone containing a Kanamycin resistance gene (GenScript Biotech) for integration into the pig genome. To prepare the plasmid for cell transfections, plasmid was grown in chemically competent E. Coli (TOP10, Thermo Fisher Scientific) and purification was done using the plasmid DNA MidiPrep kit (Qiagen) and eluted in water, following the manufacturer’s recommended protocols.

### Cell Culture

Yorkshire swine fetal fibroblasts were cultured in DMEM (Corning) with 10% (vol/vol) FBS (Atlas Biologicals) and 1x Penicillin-Streptomycin solution (Gibco). All cells were cultured in a humidified incubator at 37°C and 5% CO_2_. A Neon transfection system (Thermo Scientific) was used deliver nucleic acid or CRISPR ribonucleoprotein (RNP) to fibroblast cells. To produce fibroblast cells for somatic cell nuclear transfer, 6 x 10^5^ cells were co-transfected with two SpCas9-sgRNA RNPs, one targeting the swine *Rosa26* gene and one targeting the SRM-1 plasmid backbone, along with the SRM-1 plasmid using the 100 µl Neon transfection kit (Thermo Scientific). To prepare RNPs for transfections, lyophilized guide crRNAs and tracrRNA were resuspended in TE buffer (pH 8.0) and duplexed together by mixing 1:1 and heating to 95°C for 5 minutes before dropping to 22°C. SpCas9-gRNA ribonucleoproteins were made by incubating Cas9 protein with duplexed CRISPR gRNAs for 20 minutes at a 1:1.2 molar. After transfections, cells were plated on culture plates, and single colonies were picked for genotyping and expansion for somatic cell nuclear transfer (SCNT). To measure SRM-1 reporter activity, 6x10^4^ SRM-1 fetal fibroblasts were electroporated in triplicate with 100 ng Cre mRNA or with CRISPR RNPs using the 10 µl transfection kit (Thermo Scientific). CRISPR RNPs were prepared as described above using sgRNAs (with no crRNA/tracrRNA complexing). All reagents used for gene editing can be found in Supplemental Table 1.

### Genotyping

Fibroblast clones to be use for SCNT were screened for SRM-1 integration. Cells were grown to confluency in 96-well cell culture plates and lysed to extract genomic DNA using previously described methods^39^. The resulting lysate was PCR amplified for 35 cycles using Accustart II GelTrack PCR SuperMix (Quantabio) and run on an agarose gel to identify clones that harbored a 5’ and a 3’ integration junction amplicon. PCR primers can be found in Supplemental Table 2. Amplicons were purified using the QIAquick PCR purification kit (Qiagen) and Sanger sequencing was performed using commercial services. Clones were then analyzed for SRM-1 copy number using droplet digital PCR. For live animals, tissue biopsies were taken from the tail or ear of newborn piglets and genomic DNA was extracted using the DNeasy Blood and Tissue Kit (Qiagen) prior to running the same genotyping assays as described above.

### Somatic Cell Nuclear Transfer (SCNT) and Embryo Transfers

Fibroblast clones were shipped to TransOva Genetics for nuclear transfer using their established protocols. The resulting embryos were implanted into the uterus via midline incision and recipients were raised until farrowing or until fetuses were collected for fibroblast isolations.

### Fetal Fibroblast Isolations

Fetuses were collected 40 days into the pregnancy and skin tissue was transferred to a Type I collagenase solution at 37°C with physical agitation. Resulting cells were collected by centrifugation and cultured for 2-3 days prior to cryopreservation with FBS + 10% DMSO.

### Flow Cytometry

Fibroblast cells were lifted from tissue culture plates using TrypLE Express (Gibco) and washed with PBS before resuspension in flow cytometry buffer (1xPBS + 2.5% FBS + 5mM EDTA) to a concentration around 1x10^6^ cells ml^-1^. Flow cytometry was performed at the University of Minnesota Flow Cytometry Resource Core. Samples were analyzed utilizing a BD Biosciences LSRFortessa. Cells were analyzed for tdTomato expression, and cell viability was measured using Sytox Red (Thermo Fisher Scientific) according to the manufacturer’s recommended protocols. The resulting data files were analyzed using FACSDiva software version 9.3.

### Droplet Digital PCR

To assess SRM-1 recombination rates in fibroblast cells or from animal tissues, genomic DNA extraction was performed using the DNeasy Blood and Tissue Kit (Qiagen). DNA concentrations were measured using a Qubit 4 Fluorometer (dsDNA BR Assay Kit, Thermo Scientific). Reactions were set up using ddPCR Supermix for Probes (no dUTP) (Bio-Rad) with 20-50 ng genomic DNA per assay according to the manufacturer’s protocol. A duplexed assay was used consisting of a primer and probe set (FAM) that detects SRM-1 recombination and a diploid genomic DNA internal control primer and probe set (GAPDH, HEX). A SacI-HF restriction enzyme (NEB) was added to the reaction mix. After reaction assembly, droplets were generated using the AutoDG Instrument (Bio-Rad) followed by PCR amplification using the recommended 3-step PCR protocol with a melting temperature of 60°C. The resulting droplets were read by the QX200 Droplet Digital PCR system and subsequent analysis was performed on the QuantaSoft Analysis Pro software (Bio-Rad). SRM-1 recombination (%) was calculated as the ratio between the SRM-1 positive DNA concentration (FAM probe) and the diploid GAPDH positive DNA concentration (HEX probe) multiplied by 2 to account for the heterozygous (single-copy) SRM-1 line, then multiplied by 100 to make it a percentage. For each pig tissue sample, recombination rates were calculated and plotted using GraphPad Prism. Multiple liver and lung samples were measured and averaged for each animal in the systemic LNP study, for every other tissue only one sample was collected. To measure the SRM-1 copy number in fibroblasts for use in somatic cell nuclear transfer, the same protocol is used substituting the primer + probe (FAM) set to target the tdTomato sequence of the reporter construct. SRM-1 copy number was calculated by the QuantaSoft software, setting the GAPDH reference copies to 2. To measure the AAV-Cre copy number in pig tissues, the same protocol is used substituting the primer + probe (FAM) set to target the AAV-Cre genome sequence of the reporter construct. SRM-1 copy number was calculated by the QuantaSoft software, setting the GAPDH reference copies to 2. All primers and probes used for ddPCR in this study can be found in Supplemental Table 2.

### Animal Experiments

All swine procedures involving animals were conducted in compliance with Recombinetics IACUC and all animals euthanized were done so in accordance with AVMA approved methods. For systemic vector delivery experiments, piglets were anesthetically induced with intramuscular (IM) injection of Telazol/Ketamine/Xylazine (TKX). Anesthetic depth was verified and vitals (EKG, heart rate, temperature, respiratory rate) monitored every 10-15 minutes. Piglets were placed in dorsal recumbency and the injection site sterilely prepped. A 1-2cm incision was made to isolate the target vein where a 1.5” intravenous catheter was placed. The test article was administered over 30 seconds. The site of surgical cut-down was closed, and the animals were taken to post-operative care for observation. Upon full recovery, animals were returned to study specific group housing for in-life phases of the project. For the AAV study, AAV9 harboring a CMV enhancer and a chicken beta-actin promoter expressing Cre recombinase was purchased from the University of Massachusetts vector core (Construct AAV9.CB-PI-Cre). A litter of SRM-1 piglets was screened for neutralizing antibodies (Nab) against AAV9 in serum by the University of Pennsylvania Immunology core. An 8.8 kg male SRM-1 piglet with no detectable anti-AAV9 Nabs was injected with the AAV9 vector at a dose of 7.27 x10^12^ genome copies per kilogram body weight. A full necropsy was performed 16 days post injection and tissues were both snap frozen and fixed in 10% neutral buffered formalin overnight followed by a was in 1xPBS and transfer to 70% ethanol for further processing. For the LNP systemic study, three SRM-1 piglets were injected with 6 mL LNP carrying Cre mRNA at a dose of 0.4 milligrams mRNA per kilogram body weight (mg kg^-1^). One SRM-1 piglet underwent the same surgical procedure but was injected with 6 mL of sterile saline. Blood draws (serum and EDTA-whole blood) were performed on days 0, 1, 3, 7, and 14 on these animals as well as two sibling wild type piglets for a total of 6 samples. Samples were shipped to IDEXX Laboratories for serum chemistry and blood counts, and assay reference ranges were provided by the company. A full necropsy was performed 14 days post injection and tissues were snap frozen and fixed in 4% paraformaldehyde for further processing.

For the LNP intracerebroventricular delivery, a 3-day old piglet was placed under anesthesia and into a stereotactic injector frame. The injection site was identified through established landmarks and a Hamilton needle attached to an automated injector were used to infuse a total of dose of 80 µg of Cre-mRNA packaged in 100 µL LNP at a rate of 25 µL/min. After completion, the needle was slowly extracted, and piglet was recovered and returned to group housing for in-life phases of the study. A necropsy was performed 19 days after injection to remove the whole brain and fix in NBF for further processing.

Mouse animal procedures for LNP validation were approved by The University of California, Berkeley (UCB) institutional animal care and use committee (IACUC). All facilities used during the study period were accredited by the Association for the Assessment and Accreditation of Laboratory Animal Care International (AAALAC). To determine the *in vivo* stability and delivery efficiency of LNPs with the same formulation used for the systemic pig study, wild-type BALB/c female mice weighing between 18– 20 g were used for firefly luciferase mRNA delivery experiments. LNPs were injected into mice at a dose of 0.25 mg mRNA per kg body weight via the retro-orbital route (intravenous), and LNP/mRNA complexes were mixed with PBS pH 7.4 right before injection, resulting in a total volume of 100 μL per mouse. 4-6 hours later, a D-luciferin solution in PBS pH 7.4 solution (150 mg kg^−1^) was injected into the IP cavity of mice. After 10 min, the luciferase luminescence intensity of the whole body and organs (lungs, liver, kidneys, heart, and spleen) was imaged using an IVIS spectrum imager Figure S4. (PerkinElmer).

### *In vitro* transcription

For the ICV injection, Cre mRNA was produced by *in vitro* transcription (IVT), from a linearized plasmid template containing a CleanCap-AG-compatible T7 promoter, 5′-UTR sequence, Kozak sequence, coding sequence, 3′-UTR sequence, poly(A) tract, and a restriction site (*Esp*3I/*Bsm*BI). The 5′-UTR and 3′-UTR sequences were derived from those of Pfizer/BioNTech SARS-CoV-2 mRNA vaccine (BNT-162b2)^40^, and the Cre recombinase coding sequence was derived from the TriLink Cre mRNA open reading frame sequence (TriLink Product L-7211). Full plasmid sequence is provided in the supporting information. The IVT was performed using HiScribe T7 High Yield RNA Synthesis Kit (NEB). CleanCap Reagent AG (3′-OMe) (TriLink) was added to the reaction mixture, and UTP was replaced with N^1^-methylpseudouridine-5’-triphosphate (TriLink Biotechnologies). Following a 2-hour incubation at 37 °C, DNase I (NEB) was added to digest DNA templates. Cre mRNA was purified using the Monarch RNA Cleanup kit (NEB) and high-performance liquid chromatography.

### LNP production and characterization

For swine systemic injection studies, LNPs were formulated with an amine-to-RNA-phosphate (N:P) ratio of 4.5. The lipid nanoparticle components were dissolved in 100% ethanol with the following molar ratios: 45 mol% LP01 lipid, 44 mol% cholesterol, 9 mol% DSPC, and 2 mol% PEG2k-DMG. The mRNA cargo was dissolved in 25 mM citrate buffer (pH 4.5), resulting in a concentration of mRNA of approximately 0.45 mg/mL. LNPs were formed by microfluidic mixing of the lipid and RNA solutions using a Cytiva NanoAssemblr Spar or Ignite, following the manufacturer’s protocol, and the remaining buffer was exchanged into PBS (100-fold excess of sample volume) overnight at 4°C under gentle stirring using a 10 kDa Dialysis Cassette. The resultant mixture was then filtered using a 0.2 μm sterile filter. The filtrate was stored at 2°C-8°C. The resulting LNPs had an average size of 102.2 nm (Fig. S4). Encapsulation efficiencies were determined by the ribogreen assay^41^. Particle size and polydispersity were measured by dynamic light scattering (DLS) using a Malvern Zetasizer DLS instrument. Representative DLS spectra can be seen in Figure S3.

For ICV injection studies, LNPs were formulated following a previously published protocol^42^. A total lipid solution at a concentration of 3 mM in 100% ethanol was prepared by combining ionizable lipid 306-O12B (synthesized in house according to published protocol)^38,43^, DSPC (Avanti), cholesterol (Sigma-Aldrich), and DMG-PEG2000 (Avanti) at a molar ratio of 60:10:38.5:1.5. Cre mRNA was prepared in 10 mM sodium citrate, pH 4. The molar ratio of ionizable lipid to mRNA was 5:1. Total lipid and mRNA solutions at 3:1 volumetric ratio were combined by microfluidic mixing using the NanoAssemblr Ignite (Precision NanoSystems, Cytiva) at a 12 mL/min total flow rate. The collected mRNA-LNP sample was immediately diluted in an equal volume of 1X PBS and transferred into a Slide-a-lyzer dialysis cassette 20 kDa MWCO (ThermoFisher) for overnight dialysis in 1X PBS at 4 °C. LNP solution was filtered using a 0.22 µm syringe filter and concentrated with an Amicon 100 kDa MWCO centrifugal filter. Particle size and polydispersity index were characterized by dynamic light scattering (Zetasizer, Malvern Panalytical). Encapsulated mRNA concentration was quantified using Quant-it Ribogreen RNA Assay Kit (Invitrogen). LNP samples were kept at 4 °C until injection within 72 hours of formulation.

### Immunohistochemistry of tissues

For the AAV systemic study, complete necropsies were performed and tissues to be used for immunohistochemistry were placed in 10% neutral buffered formalin (NBF) for fixation. Small sections of tissue (1cm^3^ – 3cm^3^) were allowed to fix over a 24-hour period before being washed 3-times in PBS, larger sections (greater than 3cm^3^) allowed to fix for 72-hours, and the whole intact brain allowed to fix for 7-days. Tissues were sent to Scientific Solutions Inc. pathology service, for paraffin embedding, sectioning, and staining using their established methods. A rabbit anti-RFP antibody (Rockland 600-401-379) and a mouse anti-NIS (Abcam ab242007) were used to detect tdTomato and NIS, respectively, and scanned with a Leica Aperio CS2 whole slide scanner. Aperio ImageScope software was used to visualize and analyze the resulting digital images.

### Tissue Preparation and Fluorescent Imaging

Complete necropsies were performed and tissues to be used for fluorescent imaging were placed in 4% paraformaldehyde (PFA) for fixation. Small sections of tissue (1cm^3^ – 3cm^3^) were allowed to fix over a 24-hour period before being washed and dehydrated by transitioning to 15% then 30% of a sucrose solution in PBS. The tissues were then embedded in Tissue-Tek optimal cutting temperature compound (O.C.T., Sakura Finetek USA) and snap frozen in embedding cassettes with liquid nitrogen cooled blocks. Cryosectioning of the OTC-tissue blocks was performed on a Leica CM1860 cryostat and 5-10 µM sections were placed on positively charged microscope slides. DAPI-stain was used to highlight the nucleus of cells and coverslip adhered using Prolong Gold (Thermo Fisher Scientific). Images were captured on a Leica DM6000B microscope with an attached Leica DFC7000T microscope camera with TxRed and DAPI filters.

## Acknowledgements

We thank the entire Recombinetics laboratory and animal production staff over for their support and advice. We also thank the core facility staff at the University of Minnesota Flow Cytometry Resource, the University of Massachusetts Viral Vector Core, and the University of Pennsylvania Immunology Core as well as the staff at Scientific Solutions. Work performed at Recombinetics was supported by the Somatic Cell Genome Engineering Consortium grant from the NIH Office of the Director (U24OD026641). We thank Oleg Mirochnitchenko for his advice and guidance. Work performed at the University of Massachusetts were supported by the Rett Syndrome Research Trust. We thank Adam Hedger for synthesizing the 306-O12B lipid, and Li Li for access to the Zeta Malvern instrument to measure LNP physical characteristics. Figures were designed in BioRender.

## Contributions

JMC and DMK designed and oversaw *in vivo* studies with the help of DFC. JMC, DMK, HD, and RM ran and analyzed samples from pig studies. JMC and DAW designed and oversaw cell manufacturing and cell studies with the help of ROA. HH, SZ, and NM produced and characterized LNP-mRNA for the systemic study. YAHN and JKW produced and generated LNP-mRNA for the ICV injection study. JMC, DMK, and DFC wrote the paper with input and edits from all authors.

