## SUPPLEMENTAL DATA for "Swine reporter model for preclinical evaluation and characterization of gene delivery vectors"

### Supplementary Data

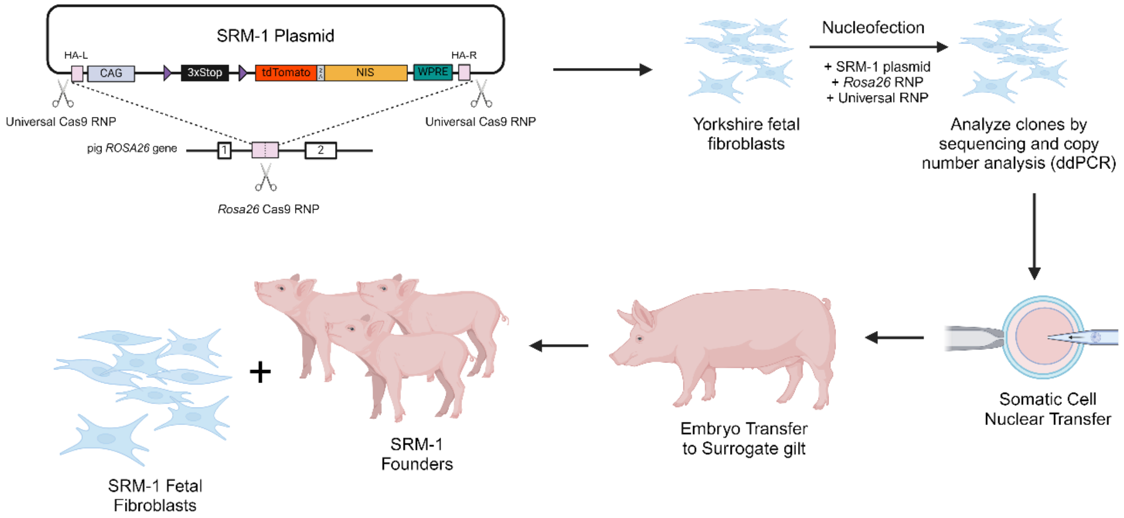


**Fig. S1 Illustration of SRM-1 model development by genome engineering and somatic cell nuclear transfer (SCNT)**. Pig cells used for somatic cell nuclear transfer were generated using genome engineering of male Yorkshire fetal fibroblast cells. The synthesized SRM-1 construct was integrated into the swine *Rosa26* gene using homology-mediated end-joining DNA repair of the SRM-1 plasmid construct after excision of the reporter from the plasmid backbone by SpCas9[44]. To stimulate integration into the *Rosa26* gene, SpCas9 RNPs targeting the first intron of *Rosa26* was co-transfected to generate a double-strand break (DSB). Short (48bp) homology arms flanking the SRM-1 reporter sequence for precise integration of the reporter by homology-directed repair. The edited cells were genotyped and cloned using SCNT. The resulting embryos were transferred to a surrogate gilt and resulting pregnancies were either carried to term to generate founder animals or collected for fetal fibroblast isolations.

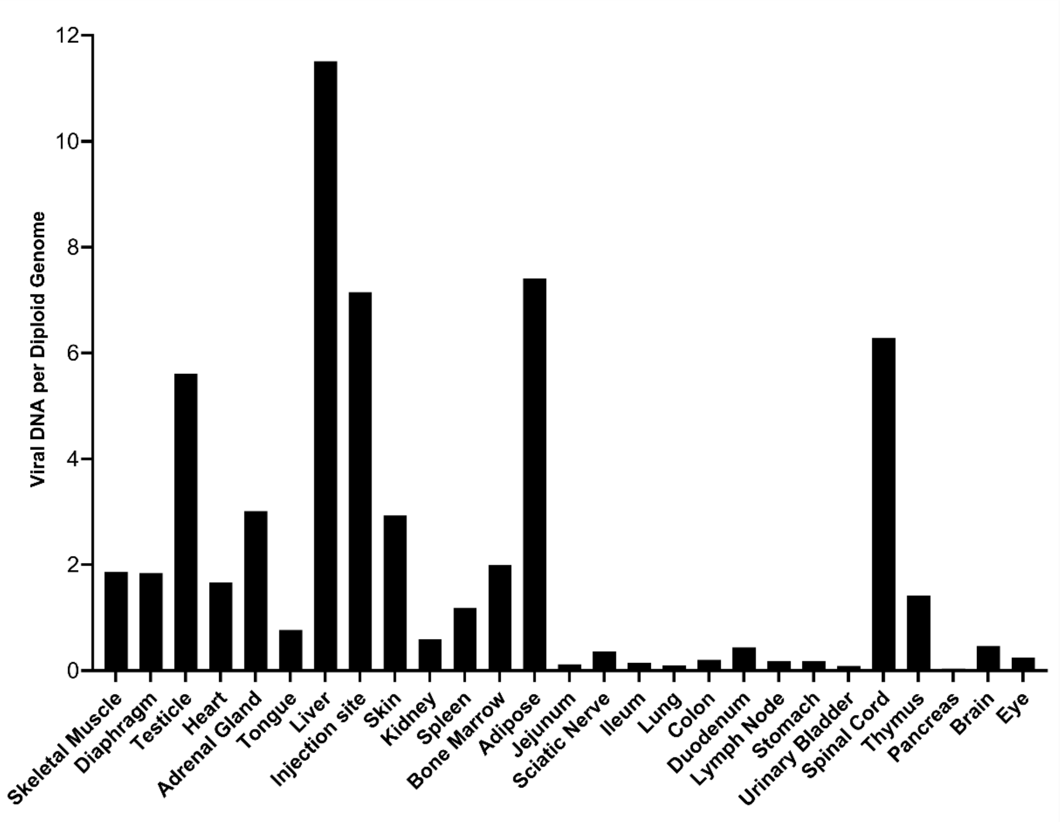


**Fig. S2 AAV9-Cre biodistribution after systemic injection.** Total DNA was isolated from each tissue in the AAV9-injected piglet and the concentration of viral DNA from the AAV genome and the concentration of a the *GAPDH* gene in pig genomic DNA were measured by ddPCR. Relative amounts of viral load in each tissue were calculated by dividing the viral DNA concentration by the diploid genomic DNA concentration.

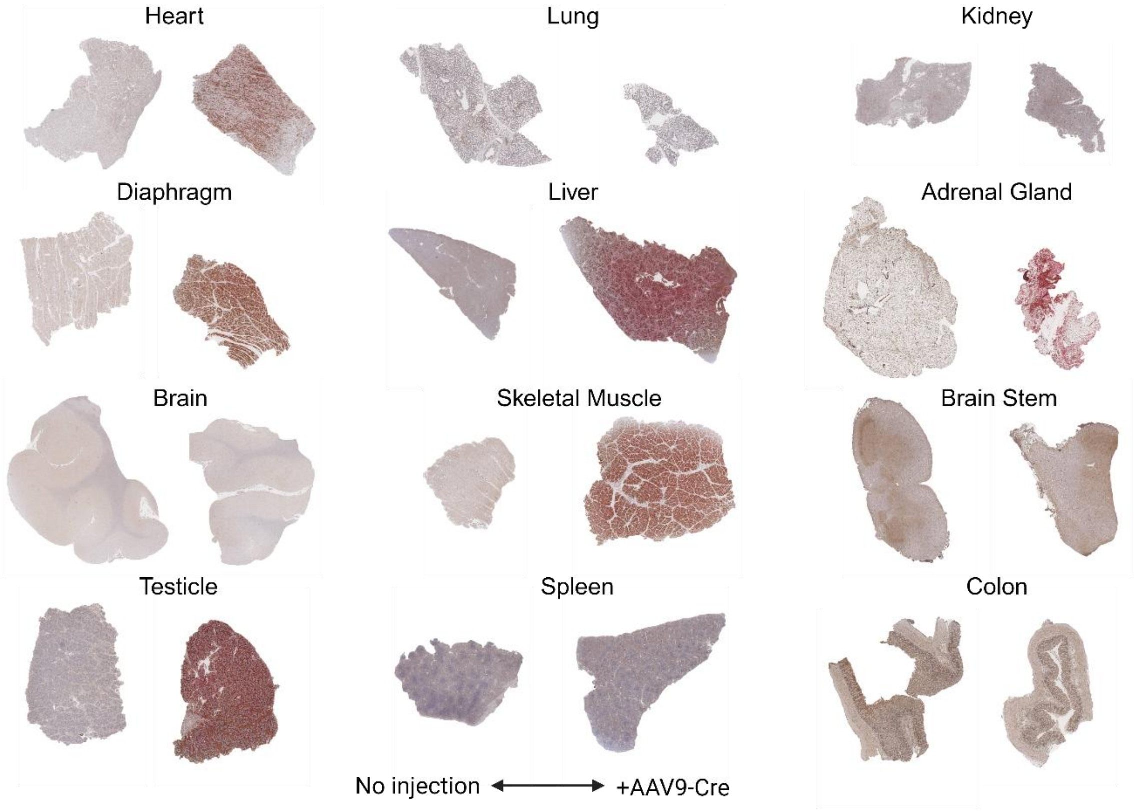


**Fig. S3 Immunohistochemistry of formalin-fixed paraffin-embedded tissue sections by tdTomato and NIS co-staining in an AAV9-Cre injected male piglet.** For each tissue, the left image shows tissues from an uninjected SRM-1 male piglet and the right image shows a male injected SRM-1 animal. The resulting tdTomato/NIS positive signal from DAB staining will be brown/red in color.

| Size/nm | PDI | EE% |
| --- | --- | --- |
| 102.2 | 0.163 | 94 |

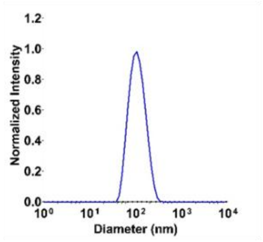


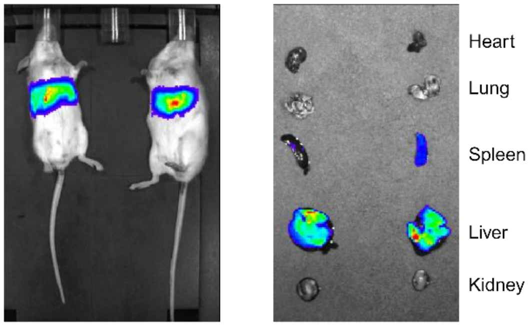


**Fig. S4 Characterization of LNP-mRNA for systemic infusions in pigs.** LNP-mRNA (Cre) size, polydisperity index (PDI), encapsulation efficiency (EE%) and an example dynamic light scattering (DLS) plot of LNPs. As a positive control, the same LNP formulation used for systemic studies in pigs was also used to make LNP-mRNA (firefly luciferase) for injection into mice. Luminescence imaging after systemic (retroobital) injection shows by in whole animal and in tissues. Highest uptake was observed in the liver and spleen.

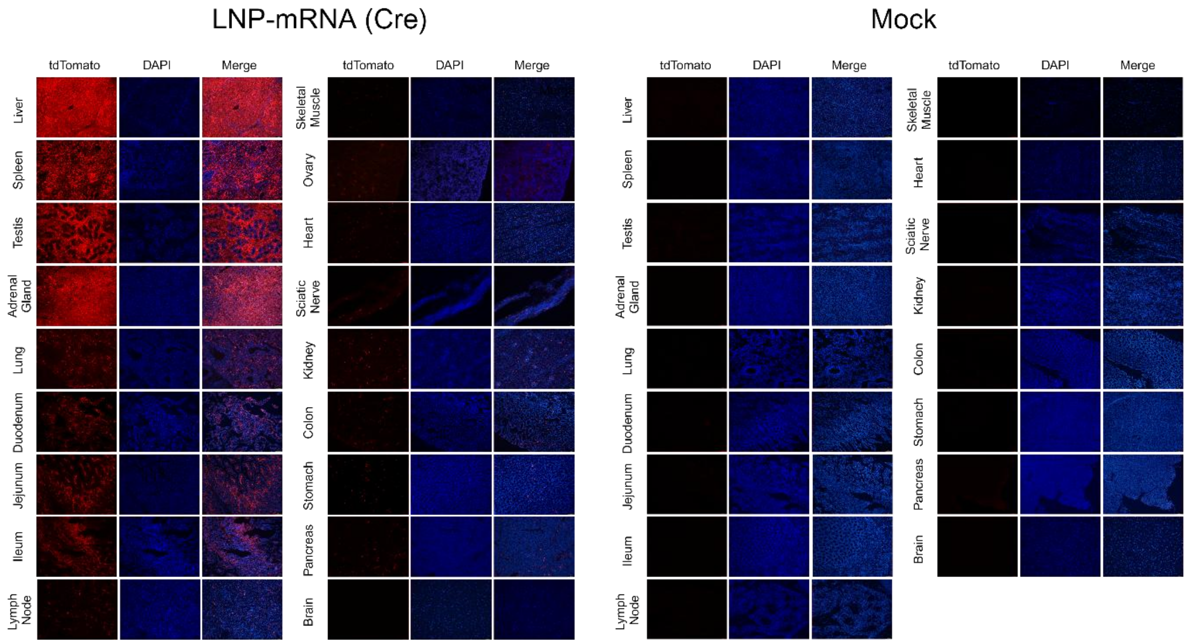


**Fig. S5 Representative fluorescent images of tissues collected from LNP-injected piglets**. Tissue samples from LNP-mRNA (Cre) injected animals (left panel), including both testicle and ovarian tissue samples from a male and female piglet, and a mock-injected male sibling animal were fixed, cryosectioned, and imaged for native tdTomato expression (red) and nuclear staining (DAPI, blue).

**Supplemental Table 1:** Synthetic sgRNAs and editing reagents

Synthetic gRNAs

| Name | Company | CRISPR Species | Spacer Sequence (5’ - 3’) | Chemical Modifications |
|---|---|---|---|---|
| ssROSA g2 | IDT | SpCas9 | GGATTTTTCTAGGCCCAGGG | AltR CRISPR-Cas9 crRNA |
| Universal gRNA | IDT | SpCas9 | GGGAGGCGTTCGGGCCACA | AltR CRISPR-Cas9 crRNA_ |
| Cas9 tracrRNA | IDT | SpCas9 | N/A | AltR CRISPR-Cas9 tracrRNA_ |
| sgRNA_sg280 | Synthego | SpCas9 | GTATGCTATACGAAGTTATT | 2'-O-Methyl at 3 first and last bases, 3' phosphorothioate bonds between first 3 and last 2 bases |
| sgRNA_sg276 | Synthego | SpCas9 | AAAGAATTGATTTGATACCG | 2'-O-Methyl at 3 first and last bases, 3' phosphorothioate bonds between first 3 and last 2 bases |
| AsCas12a | IDT | AsCas12a | GCAAAGAATTGATTTGATAC | AltR A.s.Cas12a crRNA |

Editing mRNA and proteins

| Name | Company | CRISPR Species | Modifications |
|---|---|---|---|
| Cre mRNA | TriLink BioTechnologies | N/A | CleanCap, NLS-Cre, 5-methoxyuridine modified, polyadenylated |
| AltR S.p. HiFi Cas9 Nuclease V3 | IDT | SpCas9 | Proprietary |
| AltR A.s. Cas12a (Cpf1) Ultra | IDT | AsCas12a | Proprietary |

**Supplemental Table 2:** Primers and probes used for droplet digital PCR (ddPCR) and standard PCR for agarose gel electrophoresis

Droplet Digital PCR primers and probes

| Target | Name | Fluorophore | Primer/Probe Sequence (5' - 3') |
|---|---|---|---|
| SRM-1 Cre Recombination | SRM1_ddPCR_F1 | - | CTAGAAAGTATAGGAACTTCGTCG |
| | SRM1_ddPCR_R1 | - | GCGCATGAACTCTTTGATGACC |
| | SRM1_ddPCR_Probe | FAM | CGGGATCGTGTTGCACTTAACGCG |
| SRM-1 Copy Number | ddPCR tdTomato F2 | - | GTTCCTGGGGCATGGCACC |
| | ddPCR tdTomato R2 | - | CACCTTGAAGCGCATGAACTC |
| | ddPCR tdTomato probe 2 | FAM | TGACGGCCATGTTGTTGTCCTCGGA |
| AAV9-Cre Copy Number | CreF | - | GCGGTCTGGCAGTAAAAACTATC |
| | CreR | - | GTGAAACAGCATTGCTGTCACTT |
| | Cre probe | FAM | AAACATGCTTCATCGTCGGTCCGG |
| Reference Diploid DNA | Primerpig_GAPDH_F | - | CCGCGATCTAATGTTCTCTTTC |
| | Primerpig_GAPDH_R | - | TTCACTCCGACCTTCACCAT |
| | Probepig_GAPDH | HEX | CAGCCGCGTCCCTGAGACAC |

Other PCR primers

| Target | Name | Primer Sequence (5' - 3') |
|---|---|---|
| 5' SRM-1 Junction PCR | oJMC097 | AGACTAGCTCTACCTGCTCTC |
| | PCR CMV 3-2 | GGAACTCCATATATGGGCTATGAACTA |
| 3' SRM-1 Junction PCR | LDLR HR 5-2 | GGAGGTGTGGGAGGTTTTT |
| | oJMC100 | GTAATAACACGCAGTCTCAATGC |
| SRM-1 Recombination PCR | SRM1_ddPCR_F1 | CTAGAAAGTATAGGAACTTCGTCG |
| | SRM1_ddPCR_R1 | GCGCATGAACTCTTTGATGACC |
